# An analytical framework for understanding regulatory novelty accompanying allopolyploidization

**DOI:** 10.1101/212092

**Authors:** Guanjing Hu, Jonathan F. Wendel

## Abstract

Allopolyploidy is a prevalent process in plants, having important physiological, ecological, and evolutionary consequences. Massive, genome-wide transcriptomic rewiring in response to genomic merger and doubling has been demonstrated in many allopolyploid systems, encompassing a diversity of phenomena, including homoeolog expression bias, genome dominance, expression-level dominance, and revamping of co-expression networks. Here we present an analytical framework to reconcile these patterns of regulatory novelty as governed by distinct sets of intra- and inter-subgenome *cis*-*trans* relationships. This approach is a novel extension of classic allele-specific expression analysis to incorporate and distinguish the separate effects of parental regulatory interactions as well as further complications at the allopolyploid level. We demonstrated that the *cis*-*trans* framework devised not only offers new perspective on disentangling genetic from epigenetic and higher-order effects that impact gene expression, but also provides the conceptual basis and tools to unify recently presented models for both genome-wide expression dominance and biased fractionation in allopolyploids.

---

Polyploidy, or whole-genome duplication (WGD), is exceptionally common in plants, having important physiological, ecological and evolutionary consequences (Stebbins 1940; Levin 1983; Ramsey and Schemske 2002; Leitch and Leitch 2008; Van de Peer et al. 2009; Madlung 2013; Soltis et al. 2014; Van de Peer et al. 2017; Soltis and Soltis 2016). Two types of polyploidy have long been recognized, autopolyploidy, resulting from the multiplication of one progenitor chromosome set, and allopolyploidy, involving hybridization and duplication of divergent parental genomes, classically from different species (Wendel and Doyle 2005). Allopolyploidy in particular is thought to provide avenues for regulatory novelty and hence phenotypic innovation, as evidenced by myriad non-additive and non-Mendelian responses, including gene loss and silencing (Schnable et al. 2011; Freeling et al. 2012; Liu et al. 2014; Mirzaghaderi and Mason 2017; Koh et al. 2010; Szadkowski et al. 2010; Tate et al. 2009; Anssour et al. 2009; Buggs et al. 2009; Eilam et al. 2009); activation of transposable elements (Senerchia et al. 2015; Parisod et al. 2010; Kawakami et al. 2010); epigenetic modifications (Song et al. 2017; Jackson 2017; Madlung et al. 2002; Chen 2007; Salmon et al. 2005; Rapp and Wendel 2005; Bottley 2014; Fulnecek et al. 2009; Kovarik et al. 2008; Yu et al. 2010; Shcherban et al. 2008; Wang et al. 2017; Zhao et al. 2011); and massive, genome-wide transcriptomic rewiring. The latter encompasses a diversity of phenomena, including biased expression of homoeologs on a genic (Yoo and Wendel 2014; Flagel et al. 2008; Akama et al. 2014; Combes et al. 2013; Wang et al. 2016) or even genomic (“genome dominance”) scale (Edger et al. 2017; Yang et al. 2016; Zhang et al. 2015; Flagel and Wendel 2010; Schnable et al. 2011; Garsmeur et al. 2014); the poorly understood phenomenon of “expression level dominance” (Rapp et al. 2009; Yoo et al. 2013; Grover et al. 2012; Zhang et al. 2016; Akhunova et al. 2010; Liu et al. 2014); and the modification of duplicated gene co-expression networks (Gallagher et al. 2016; Hu et al. 2016). A hallmark of these phenomena is deviation from vertical transmission of preexisting patterns, or the “parental legacy”, inherited from the two progenitors (Buggs et al. 2014). These deviations collectively represent regulatory novelty that either accompanied or evolved following genome merger and doubling.

Notwithstanding this progress in our understanding of expression alteration accompanying allopolyploidization, there remains a need to develop further an encompassing conceptual framework. Here we propose such a framework based on the characterization of regulatory divergence between parental species and subsequent changes at the allopolyploid level. In terms of parental divergence, identifying the types of regulatory changes that have evolved between diploids has long been a focus of classical allele-specific expression (ASE) analysis (Wittkopp et al. 2004). That is, allele-specific expression in F1 hybrids provides a readout of relative *cis*-acting activity in a common *trans* environment, whereas expression differences between parental species not attributed to *cis*-acting divergence are inferred to be caused by *trans*-acting variations. However, how interactions among divergent regulatory alleles affect gene expression in natural hybrid and allopolyploid species remain poorly understood, so uncovering the interplay between the ever-evolving *cis* and *trans* elements is critical for understanding phenotypic innovations that emerge following genomic merger and doubling. Here we extend the classical ASE model to the polyploid level, by considering the duplicated sets of *cis*-*trans* relationships initiated by interspecific hybridization.

As illustrated in Figure 1A, we use the cotton (*Gossypium* L.) allopolyploid system as an example, as it is illustrative of many of the model systems used today in studies of polyploidy. Allotetraploid (“AD genome”) cottons originated ~1-2 million years ago from a hybridization event between two diploid species (“A” and “D”) followed by whole-genome duplication (Wendel and Grover 2015; Wendel and Cronn 2003; Wendel et al. 2010). The descendants of the parental diploid species remain extant (“A2” and “D5”), from which a synthetic F1 hybrid was generated; this has been used to disentangle expression changes due to hybridization from those arising later from polyploidy and subsequent evolution (Yoo et al. 2014; Flagel et al. 2008; Flagel and Wendel 2010). For the synthetic F1 hybrid and natural tetraploid cottons, the expression of each pair of duplicated genes (homoeologs “At” and “Dt”, with “t” denoting subgenome) is governed by four sets of *cis*-*trans* relationships, including two intra-subgenome interactions derived from each of the parental diploids (*aa* and *dd*), and two newly formed inter-subgenome interactions (*ad* and *da*).

**Figure 1.**
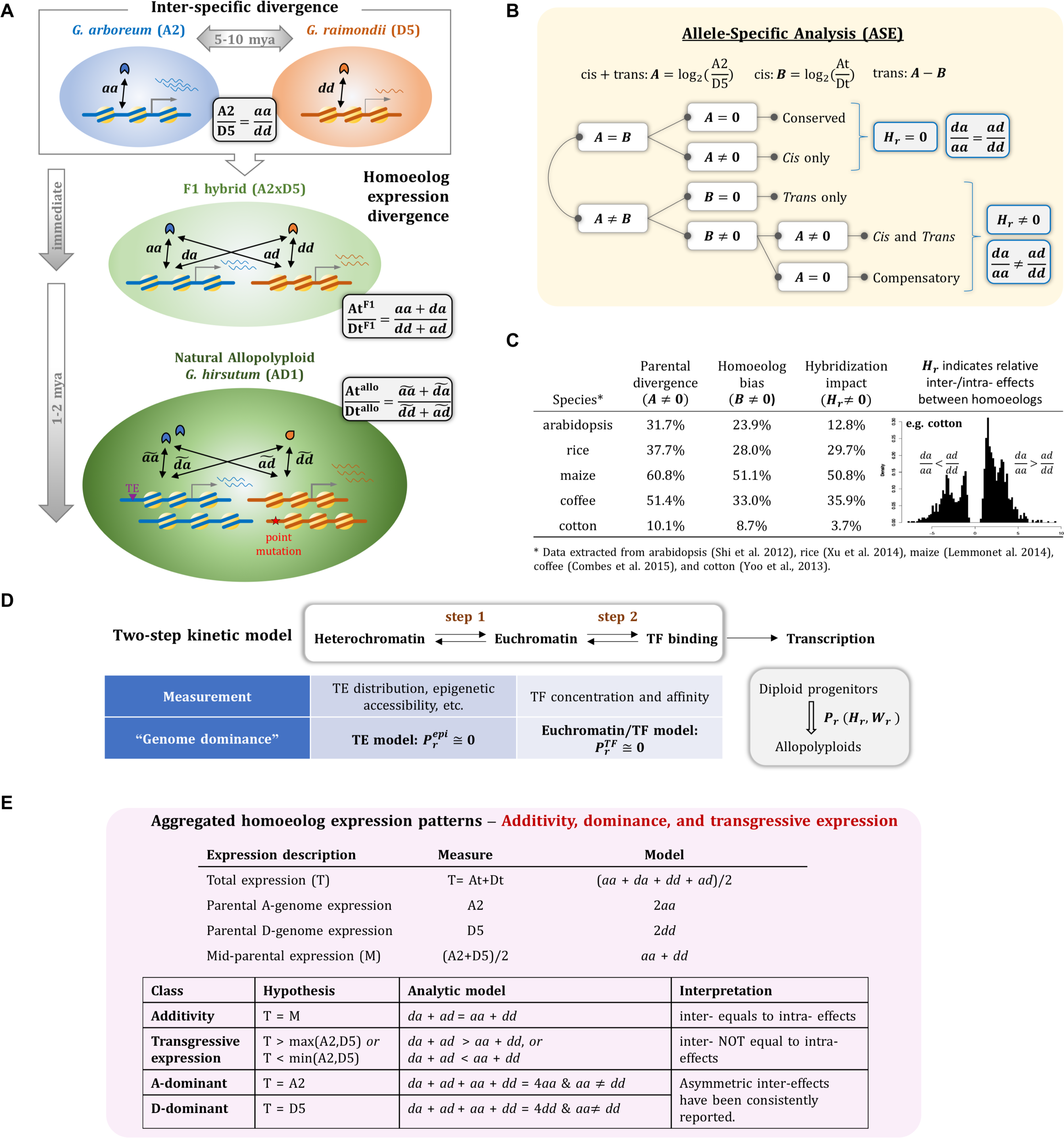
A proposed analytical framework for understanding regulatory novelty accompanying hybridization and allopolyploidy, using the cotton (*Gossypium* L.) allopolyploid system as an example. (**A**) Between the parental diploid species *G. arboreum* (A2) and *G. raimondii* (D5), differential gene expression and/or chromatin accessibility are determined by the divergence of corresponding intra-genome *cis*-*trans* interactions *aa* and *dd*, respectively. Following genomic merger and doubling, the At and Dt homoeolog divergence is governed by two more sets of newly formed inter-subgenome interactions *ad* and *da* (first letter indicates *trans* origin, and second letter indicates *cis* origin). In natural allopolyploids, stoichiometric changes and sequence evolution (e.g. TE insertion and point mutation) may further alter *cis*-*trans* interactions to become 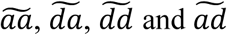. (**B**) A schematic diagram of classic allele-specific expression analysis (ASE), with interpretations based on relative inter-versus intra-*cis*-*trans* interactions noted in blue boxes. (**C**) Percentages of parental divergence, homoeolog expression bias and hybridization impact (*H*_*r*_) in various plant systems. The histogram of significant *H*_*r*_ is shown for cotton. (**D**) According to a two-step kinetic model of gene transcription, two hypotheses of “genome dominance”, the euchromatin/TF model (Bottani et al. 2018) and the TE model (Wendel et al. 2018; Steige and Slotte 2016), are complementary to each other. These provide a conceptual framework for revealing the mechanisms that underlie novel *cis*-*trans* interactions. (**E**) Understanding non-additive expression patterns as regulated by *cis*-*trans* interactions.

According to the ASE model (Wittkopp et al. 2004), regulatory divergence acting only in *cis* between the parental diploids will be mirrored as allele-specific expression in the hybrid and polyploid (At/Dt = A2/D5, where At, Dt, A2 and D5 refer to expression levels for those genic copies; Figure 1B). Any deviations from the parental divergence (i.e., At/Dt ≠ A2/D5) can be assigned to the influence of *trans* variation, either acting only in *trans* (At/Dt = 1, because of the common *trans* environment) or by variants acting both in *cis* and *trans* (At/Dt ≠ 1). The latter combinatorial effect may also be invisible prior to interspecific hybridization (A2/D5 =1 and At/Dt ≠ 1), as *cis* and *trans* variants may be compensatory. Such “compensatory” patterns have been suggested to result from stabilizing selection in order to maintain conserved gene expression during parental divergence (Shi et al. 2012; Tirosh et al. 2009), whereas cross interactions between the independently derived genetic variants give rise to immediate expression novelty following genomic merger. In comparison with the “*cis* only” and “*trans* only” effects that have been extensively studied in plant hybrid and allopolyploid systems (Shi et al. 2012; Lemmon et al. 2014; Xu et al. 2014; Combes et al. 2015; Chaudhary et al. 2009; Springer and Stupar 2007; Bell et al. 2013; He et al. 2016), understanding the molecular basis of the combinatorial effects of *cis* and *trans* variants has been challenging. For example, did the co-evolution of *cis* and *trans* elements occur in one or both diploid species? How did they evolve from their ancestral states? How do “foreign” (those from different genomes or species) and “native” (from the same genome or species) interactions differ? Is it possible and/or to what extent are “native” and “foreign” interactions preferred? That is, are these relationships asymmetrical? The common *trans* environment shared by both homoeologs has made these questions conceptually difficult to investigate.

Therefore, let us consider more specifically how *trans* regulators of different origins act on their cognate and cross-genome targets. For instance, expression of the At homoeolog is determined through its own *cis* elements interacting with both the A- and D-genome *trans* factors (represented by *aa*+*da*), while expression in the diploid parent is attributed to only the *cis*-*trans* relationship native to the A-genome diploid (*aa*). Thus, the contrast between the homoeolog-specific expression (At/Dt) and parental expression divergence (A2/D5) can be modeled as:

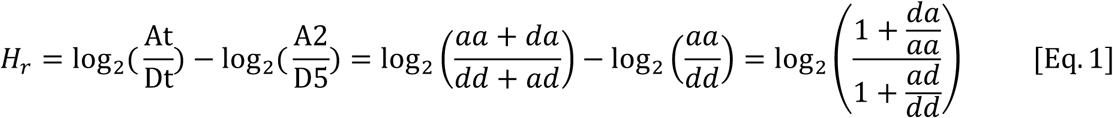

where *H*_*r*_ represents the impact of hybridization on relative homoeolog expression, and is also equivalent to the additive inverse of the *trans* effect 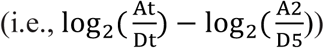 estimated in classic ASE analysis (Figure 1B). This acknowledges that hybridization inherently affects homoeolog-specific expression in *trans*, dependent on the relative effects of inter-versus intra-subgenome interactions. Although the foregoing algebraic inference is not substantially different from that of classic ASE analysis, the perspective is nonetheless meaningful. Not only is the impact of hybridization, *H*_*r*_, conceptually distinguished from how *cis* and *trans* variants contribute to parental divergence, but Eq. 1 also presents a method to quantify how inter-subgenome interactions differentially regulate each homoeolog relative to intra-subgenome interactions (*da/aa* vs. *ad/dd*). As summarized in Figure 1C, the impact of hybridization varies across plant systems, and correlates with the amount of expression divergence between parental species. A histogram of significant *H*_*r*_, as exemplified for cotton (Yoo et al. 2013), is indicative of asymmetrical regulation by cross-genome interactions; that is, inter-subgenome interactions have a stronger relative effect on At than Dt. This realization focuses attention on inter-subgenome interactions, which are most relevant to gene expression alteration accompanying hybridization *per se*.

In comparison with the *trans* action of hybridization *per se*, how genome *doubling* alters homoeolog gene expression is complicated by multiple issues of scaling and stoichiometry. With the increase of DNA content accompanying allopolyploidy, imperfect proportionalities and non-linear relationships with cellular and nuclear volumes set in motion a cascade of stoichiometric imbalances (among, for example, transcriptional machineries and transcription factors), which collectively alter gene expression. Because the physiochemical responses of individual homoeologs vary from gene to gene, it is not yet possible to systematically predict how stoichiometric imbalances triggered by genome merger and doubling will impact regulatory interactions. It does appear, however, that the increased range of homoeolog-specific expression (when the variance of 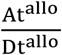 is larger than 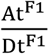) is expected, as reported in cotton (Yoo et al. 2013), wheat (Wang et al. 2016) and rice (Xu et al. 2014; Sun et al. 2017).

In a *cis-trans* framework, genome doubling can be modeled by contrasting homoeolog-specific expression between the corresponding allopolyploid and the F1 hybrid. The impact of genome doubling *W*_*r*_ is as follows:

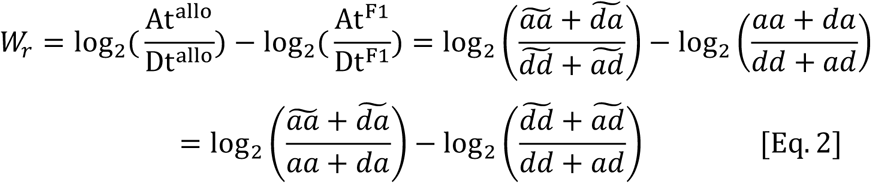

where the *cis*-*trans* interactions in allopolyploids are marked with accent as 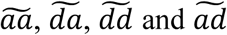. The same notion applies to the overall effect of allopolyploidization, *P*_*r*_:

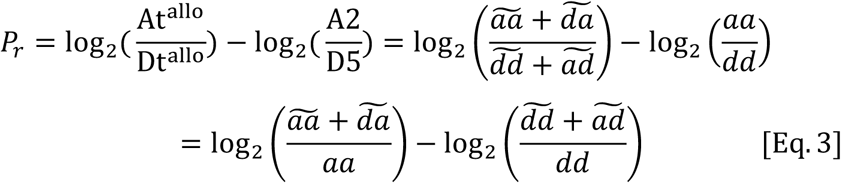

Thus, the emergence of polyploid-specific patterns (*W*_*r*_ ≠ 0 and/or *P*_*r*_ ≠ 0) depends on the alteration of any or all *cis*-*trans* interactions, which in natural allopolyploids will ensue from a spectrum of stoichiometric responses, epigenetic remodeling and genetic changes.

How these changes collectively affect regulatory interactions are relevant to several of the principal generalizations about gene expression in allopolyploids. For example, under what circumstances do these interactions preferentially shift homoeolog expression ratios towards one progenitor or the other (e.g. 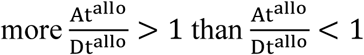)? In other words, how might altered *cis*-*trans* interactions in allopolyploids account for “genome dominance” (Schnable et al. 2011)? Similarly, how might this perspective shed light on the observation of preferential or biased transcription of one of the two co-resident genomes in an allopolyploid (“unbalanced homoeolog expression bias” at the genomic scale (Grover et al. 2012))?

One possible insight into the mechanisms underlying these dynamics is offered by Bottani et al. (2018), who adapted a kinetics model of transcription factor (TF) binding (Bost and Veitia 2014; Chu et al. 2009) to allopolyploidy. They demonstrated that when cross-regulation is involved, parental difference in TF affinity rather than concentration is the key driver of differential transcription response for homoeologous target genes. Thus, a dominant effect of the parental TF that displays higher affinity presents a quantitative explanation for “gene dominance” immediately after allopolyploidy. Next, the authors expanded the simple one-step model of TF binding to two sequential reactions – first the establishment of chromatin accessibility, then transcription factor binding to the accessible promoter site to activate transcription (Figure 1D). This additional epigenetic component thereby provides a testing ground to extend gene-level dominance to the genomic/epigenomic scale. For example, for the parental genome with a larger euchromatic content, more nonfunctional binding sites are accessible, hence creating a higher noise/signal level from the perspective of TF binding. Thus, TF binding affinities and/or concentrations are expected to scale with accessible euchromatin content, all else being equal, in which case, higher TF binding affinity is a likely outcome. As Bottani et al. (2018) showed, when homoeologous TF affinities are proportional to the ratio of euchromatic content, the parental genome with larger euchromatic content is destined to display higher transcription activities following genomic merger and doubling, hence becoming the “dominant” subgenome in allopolyploids.

It is worth nothing that this quantitative model is, to some extent, in line with the prevailing explanation for biased homoeolog expression and biased genome fractionation, based on the “genomic legacy” of TE contents (Wendel et al. 2018; Steige and Slotte 2016). This explanation has been conceptualized in recent years from accumulating literature on chromatin modification, TE content, and small RNA biology (Springer et al. 2016; Diez et al. 2014; Zhang et al. 2017; Renny-Byfield et al. 2017; Yang et al. 2016). Phrased simply, the different parental states of TE load and distribution between sub-genomes lead to differentiated epigenetic control (e.g. small RNA populations and preferential recruitment of epigenetic modifiers) for homoeolog expression, and as a consequence the homoeolog physically closer to epigenetically silenced TEs is more likely to be repressed via localized heterochromatinization, and even lost in the longer term (hence, “biased fractionation”; see recent review by Wendel et al. (2018)).

A key difference between the euchromatin/TF model (Bottani et al. 2018) and the TE model (Wendel et al. 2018; Steige and Slotte 2016), which also makes them complementary to each other, is that the euchromatin/TF model is dependent on a difference in TF affinities and euchromatin content that co-evolved in parental genomes, whereas the TE/biased-fractionation model mainly considers differences in chromatin accessibility and gene expression mediated by parental TE adjacency (Figure 1D). What the two models share is the requirement of inheritance of differentiated parental conditions, one being TF affinity while the other is TE adjacency. By analogy to studying the impact of allopolyploidy on homoeolog expression ratios (*P*_*r*_), as defined above, the effects of inheritance of these parental states can be tested by, with superscripts denoting partitioning of mechanistic effects, 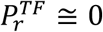 for the measure of TF affinity, and 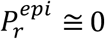 for TE adjacency and/or epigenetic accessibility. In reality, both scenarios are likely to be intertwined in natural situations, and may even be in conflict with each other. For example, two homoeologs may differ in terms of regulator TF affinity (for whatever reason), but the homoeolog with stronger TF binding may still be expressed at a lower level due to a nearby TE insertion. On the other hand, two homoeologs that differ in promoter accessibility may still be equally expressed, if stronger TF affinity is newly gained for the less accessible homoeolog, or the less accessible promoter has gained more functional binding sites since allopolyploidy. Obviously, a co-examination of both scenarios is most likely to uncover the determinative mechanisms for homoeolog expression divergence.

The foregoing considerations present both a useful *cis-trans* framework and an experimental agenda for disentangling genetic from epigenetic and higher-order effects that impact gene expression in hybrids and allopolyploids. Of course, translating this conceptual structure into empirical estimates will require more than just expression data, but access to genetic and epigenetic regulatory information is now within reach in many systems. For example, a spectrum of technologies is available to interrogate transcription factor binding to promoters (Bartlett et al. 2017; Jin et al. 2017; Landt et al. 2012; Weirauch et al. 2014), and similarly, a range of chromatin assays (Lane et al. 2014; Celniker et al. 2009; Lu et al. 2017; Jiang 2015; Zentner and Henikoff 2012) are now practical that permit the assessment of the relative accessibility of homoeologs and orthologs to the transcriptional machinery; one was recently applied in maize to connect chromatin states with biased fractionation following ancient polyploid event (Renny-Byfield et al. 2017). It is the joint application of these technologies with expression data, using the conceptual partitioning described here, which will facilitate new understanding of duplicate gene behavior in hybrids and polyploids.

In addition to homoeolog-specific expression patterns of expression bias and genome dominance, several key phenomena that characterize novel patterns of aggregated homoeolog expression have also been extensively studied, such as additive and non-additive expression, expression-level dominance, and transgressive expression, as reviewed (Yoo et al. 2014). Interpreting these patterns across systems remains an issue due to terminological inconsistency (Grover et al. 2012), among other factors. Perhaps more germane is the point that conceptual relationships among these different phenomena are not well understood, thereby impeding the synthesis required to uncover the underpinnings of duplicate gene expression evolution. The approach outlined here may facilitate such an understanding, by focusing attention on the interplay between genomic legacy features such as TE adjacency and chromatin state, biophysical interactions such as TF binding efficiency, and how these ancestral as well as newly formed *cis*-*trans* relationships govern expression evolution accompanying genome merger and doubling. As examples, we highlight two broad questions for which our conceptual framework may find utility:

1. **To what extent do homoeolog expression bias and non-additivity reflect novel, *cis/trans* interactions?** Homoeolog expression bias is when one of two duplicated genes (homoeologs) is expressed more than the other; that is, 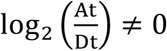. As modeled in Figure 1A, four sets of inter-and intra-subgenome interactions are involved, and even the parental sets may have been altered following genomic merger and doubling (i.e., 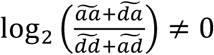). The amount of homoeolog expression bias that resembles parental divergence is relatively consistent among plant species (under 20%), whereas the amount of expression bias attributed to cross-genome interactions and other types of alterations is more variable (1.4%-37.8%); these estimates were extracted from studies of widely diverged plants - arabidopsis (Shi et al. 2012), cotton (Yoo et al. 2013), maize (Lemmon et al. 2014), rice (Xu et al. 2014) and coffee (Combes et al. 2015). Similarly, to test for expression additivity, it is common to compare *total* expression for a pair of homoeologs (T = At + Dt) to the average of parental expression values 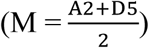. Because current methods like RNA-seq rely on per-transcriptome normalization to compare expression level across samples, there is an underlying assumption of equal *transcriptome size* (Visger et al. 2017); this, however, likely is not true in most cases (Coate and Doyle 2010) due to the multiple stoichiometric and volumetric cascades that affect gene expression following hybridization and doubling. As shown in Figure 1E, additive expression patterns are determined by the equal *total* inter- and intra-effects, which has no direct equivalence with any ASE category (Figure 1B). Non-additive expression patterns, including expression-level dominance and transgressive expression levels, arise from all four sets of regulatory interactions, these reflecting complex non-linear biochemical and biophysical interactions. This may help explain the large variation in non-additive expression patterns, ranging from less than 1% to 7% in different allohexaploid wheat species (Chelaifa et al. 2013; Chague et al. 2010), from 23 to 61% among variable cotton tissues (Rambani et al. 2014; Flagel and Wendel 2010; Yoo et al. 2013), and from 42% to 60% under two temperature conditions in coffee (Bardil et al. 2011). Teasing apart the mechanistic basis of these novel *cis/trans* interactions poses an interesting research challenge for future studies.
2. **How is the direction of expression level dominance determined by *cis* and *trans* regulation?** It has been suggested that expression-level dominance toward one parent is mainly caused by up- or down-regulation of the homoeolog of the “less dominant” parent (Shi et al. 2012; Yoo et al. 2013; Cox et al. 2014; Combes et al. 2015). Taking the A-dominant expression pattern as an example (Figure 1E, see “A-dominant” row), the joint effect of inter- and intra-interactions approximates the effect of equal number of intra A-genome regulation; if up- or down-regulation is mainly observed for the Dt homoeolog to approach the A-like expression (i.e., *dd* + *ad* = 2*aa*), the intra- and inter-effects of At are equal to each other (i.e., *da* = *aa*). This implies that the At expression is mainly determined by its *cis* element regardless of the origin of *trans* factors, while at the same time the Dt expression is under strong influence of the At *trans* factors. Thus, expression level dominance is likely to be associated with divergent *trans* factors between diploid progenitors, and the progenitor with stronger, more influential *trans* factors will become dominant with respect to *total* gene expression. In this context, it will be interesting to explore whether candidate *trans* factors such as TFs are differentiated between homoeologs in terms of concentrations and affinities. It will also be interesting to evaluate whether the strong *cis* effect of the dominant homoeolog is caused by sequence variation of binding motifs or by a more accessible chromatin formation. Because inter-subgenome interactions can up- or down-regulate target homoeologs, the direction of expression level dominance appears *not* to be associated with the direction of homoeolog bias; it will be interesting to parse the underlying mechanisms of this distinction.

Beyond the gene-centric characterization of expression changes, another relevant and pressing question concerns how gene-to-gene networks are reshaped by genomic merger and doubling, in terms of the genome-wide collection of inter- and intra-subgenome interactions? As recently reviewed by Gallagher et al. (2016), co-expression network analysis in polyploids not only has the potential to facilitate a better understanding of the complex ‘omics’ underpinnings of phenotypic and ecological traits, but also may provide novel insight into the interaction among duplicated genes and genomes. Given that previous work in allopolyploids (e.g. wheat (Pfeifer et al. 2014) and cotton (Hu et al. 2016)) are mainly based on aggregated co-expression relationships of homoeologs, one future direction is to generate networks considering homoeolog expression separately, thereby allowing the direct evaluation of topological dynamics in terms of gain and loss of intra- and inter-subgenome relationships (Conant 2010; Conant and Wolfe 2008; Conant and Wolfe 2006). Although co-expression relationships do not necessarily represent physical interactions between *cis* and *trans* regulatory elements, the gene-to-gene interconnections that are inferred based on the “guilt-by-association” principle provide an alternative, and parallel approach to estimate the impact of genomic merger and doubling, under the same analytic framework as proposed. Future analyses of gene networks could include integration with parental *cis*-*trans* divergence, novel cross-genome interactions, and various expression-level phenomena, together with other epigenetic and physiochemical datasets.

In conclusion, the opportunity to advance our understanding of transcriptome dynamics in hybrids and allopolyploids is being enabled by the maturation of multiple “omics” technologies and conceptual advances, the latter including a focus on the mechanistic underpinnings of intergenomic *cis*-*trans* interactions, as explicated here. It is likely that these perspectives and approaches will yield new insight into the origin of physiological and phenotypic responses to hybridization and polyploidy, and thereby to the evolutionary process in general.

## Acknowledgements

We thank Corrinne E. Grover and Justin Conover for helpful discussion and comments on the manuscript. We are also grateful for comments and criticisms from the reviewers and editors, which prompted us to revise and substantially improve this manuscript. Research in the Wendel laboratory has been funded by NSF, whose support we gratefully acknowledge.

